# Test-retest reliability of spectral parameterization by 1/*f* characterization using *SpecParam*

**DOI:** 10.1101/2023.09.20.558566

**Authors:** Daniel J. McKeown, Anna J. Finley, Nicholas J. Kelley, James F. Cavanagh, Hannah A. D. Keage, Oliver Baumann, Victor R. Schinazi, Ahmed A. Moustafa, Douglas J Angus

**Author notes:** **Corresponding author:** Dr. Daniel J. McKeown Bond University, Gold Coast Queensland 4229, Australia. ORCiD: 0000-0003-0298-6806.

## Abstract

*SpecParam* (formally known as *FOOOF*) allows for the refined measurements of electroencephalography periodic and aperiodic activity, and potentially provides a non-invasive measurement of excitation:inhibition balance. However, little is known about the psychometric properties of this technique. This is integral for understanding the usefulness of *SpecParam* as a tool to determine differences in measurements of cognitive function, and electroencephalography activity. We used intraclass correlation coefficients (ICC) to examine the test-retest reliability of parameterized activity across three sessions (90 minutes apart and 30 days later) in 49 healthy young adults at rest with eyes open (EO), eyes closed (EC), and during three EC cognitive tasks including subtraction (Math), music recall (Music), and episodic memory (Memory). ICCs were good for the aperiodic exponent and offset (ICCs > .70) and parameterized periodic activity (ICCs > .66 for alpha and beta power, central frequency, and bandwidth) across conditions. Across all three sessions, *SpecParam* performed poorly in EO (40% of participants had poor fits over non-central sites) and had poor test-retest reliability for parameterized periodic activity. *SpecParam* mostly provides reliable metrics of individual differences in parameterized neural activity. More work is needed to understand the suitability of EO resting data for parameterization using *SpecParam*.

## INTRODUCTION

The electroencephalogram (EEG) is characterized by fluctuations in both periodic (oscillatory) and aperiodic (non-oscillatory) activity (Keil et al. 2022; Wen and Liu 2016). The development of novel techniques has allowed researchers to independently parameterize periodic and aperiodic activity. This has produced new insights into intrinsic and task-related brain function. Although total periodic activity has been studied extensively (Basar 2013; Klimesch 1997; 1999), aperiodic activity has only recently been recognized as physiologically and mechanistically valuable for understanding brain function (Keil et al. 2022; McNab et al. 2015; Voytek et al. 2015). Both periodic and aperiodic activity vary with cognition (Cross et al. 2022; Finley et al. 2023; Immink et al. 2021; Thuwal et al. 2021a), disease states (Kim et al. 2022; McKeown et al. 2023; Rosenblum et al. 2023; Wang et al. 2022c), developmental disorders (Arnett et al. 2022; Karalunas et al. 2022; Ostlund et al. 2021; Shuffrey et al. 2022), and across the lifespan (Brady and Bardouille 2022; Finley et al. 2022; Hill et al. 2022; Merkin et al. 2023; Thuwal et al. 2021b).

All techniques and measurement instruments require good psychometrics to be useful for understanding individual or group differences, yet little is known about the psychometric properties of EEG following parameterization techniques. As measures of EEG activity in the absence of parameterization (i.e., total periodic activity; Gasser et al. 1985; Popov et al. 2023) has traditionally been employed to assess psychometric properties of EEG, only recently the psychometric properties of periodic activity been assessed following parameterization (i.e., parameterized periodic activity; Levin et al. 2020; Pathania et al. 2021) while also focusing on aperiodic activity (Levin et al. 2020; Pathania et al. 2021; Popov et al. 2023; Trondle et al. 2023; Webb et al. 2023). This is integral to our understanding of EEG correlates of neural activity, as aperiodic activity conflates all periodic activity in each frequency band. Therefore, in the current study, we extend research examining the test-retest properties of parameterized periodic and aperiodic activity and offer caveats and considerations for their use as biomarkers.

### Periodic and Aperiodic Activity

Although oscillatory activity within canonical frequency bands (delta 1 – 4 Hz, theta 4 -7 Hz, alpha 7 -13 Hz, beta 13 – 30 Hz, and gamma >30 Hz) has typically been the focus of studies examining spontaneous and task-driven neural activity, the EEG signal is predominantly made up of non-oscillatory aperiodic activity (Donoghue et al. 2022). Aperiodic activity is present across all frequencies and follows a 1/*f* distribution, where power decreases as a function of frequency. Because periodic and aperiodic activity co-occur and may change independently, the systematic variation in aperiodic activity can create the appearance of systematic variation in total periodic activity (Donoghue et al. 2020b). That is, changes and individual differences in total periodic activity, which traditionally involves quantifying the total power in the canonical frequency range irrespective of the aperiodic component, can be conflated with changes and individual differences in aperiodic activity. Indeed, in a recent study assessing changes in the theta-beta ratio across adulthood (Finley et al. 2022), changes in the ratio were highly correlated with the aperiodic component (*r* = .71).

New methods (Donoghue et al. 2020b) allow for the independent quantification of parameterized periodic (e.g., parameterized theta, alpha, and beta) and aperiodic parameters (e.g., offset, and exponent). The *SpecParam* algorithm (Donoghue et al. 2020b), formally known as *Fitting Oscillations and One Over f* (*FOOOF*), isolates the parameterized periodic activity presented as peaks on the power spectral density (PSD) after the ’removal’ of the aperiodic component. This allows for the quantification of three parameterized periodic activity measures: power, central frequency, and bandwidth. Power is the height above the aperiodic ‘baseline’ activity, central frequency is the most prominent part of the periodic peak, and bandwidth is the range of frequency around the central frequency of the peak which shows elevated periodic activity. Other formal methods, such as *Pink and White Noise Extractor* (*PaWNextra*; Barry & De Blasio, 2021), *Multiple Oscillation Detection Algorithm* (*MODAL*; Watrous et al., 2018), *Extended Better Oscillation Detection* (*eBOSC*; Kosciessa et al., 2020), and *Irregular-Resampling Auto-Spectral Analysis*’ (*IRASA*; Wen & Liu, 2016) are less frequently used, and either focus on the distinction between pink (1/*f*-like) and white noise (*PaWNextra*) in PSDs, or extract aperiodic activity directly from the processed MEG/EEG signal (*MODAL*, *eBOSC*, and *IRASA*).

The slope of the PSD (i.e., the aperiodic exponent) has been causally linked to excitation:inhibition (E:I) balance (Gao et al. 2017; Muthukumaraswamy and Liley 2018). Recently, Lendner et al. (2023) demonstrated in a series of experiments that the aperiodic slope in the human EEG is similar to the aperiodic slope in the rodent scalp EEG, which corresponds to changes in pyramidal neuron (excitation) and interneuron (inhibition) activity derived from two-photon calcium imaging (Grienberger et al. 2022). That is, the slope increased as pyramidal neuronal activity was high (increased excitability) and decreased when pyramidal neurone activity was higher than inhibitory interneuron activity. Consistent with basic and translational neuroscience findings (He 2014; Voytek and Knight 2015), studies in human participants have found that variation in the aperiodic exponent is associated with cognitive state (Donoghue et al. 2020a), disease states (McKeown et al. 2023; Wang et al. 2022c), developmental disorders (Ostlund et al. 2021), age (Finley et al. 2022; Merkin et al. 2023), and cognitive decline (Finley et al. 2023). Steeper slopes – presumably reflecting a relative reduction in E:I balance – have been observed in conditions associated with an increase in inhibitory neural activity (e.g., propofol anesthetic; Waschke et al. 2021) and in individuals characterized by relatively greater inhibitory to excitatory activity (e.g., younger adults compared to older adults; Finley et al. 2022; Merkin et al. 2023). Conversely, flatter slopes – presumably reflecting a relative increase in E:I balance – have been observed in conditions that require the rapid inhibition of neural activity (e.g., cognitively taxing tasks), and in populations with neural hyperexcitability, such as adolescents with Attention-Deficit Hyperactivity Disorder (ADHD; Ostlund et al. 2021). Although less frequently examined, several studies have found that the height of the PSD (i.e., the aperiodic offset) is associated with disease states such as Parkinson’s Disease (McKeown et al. 2023), and is predictive of better performance in reactive control tasks (Clements et al. 2021). These findings are broadly consistent with observations that the aperiodic offset is coupled to overall neural spiking rates and is strongly correlated with blood-oxygen level-dependent (BOLD) activity (Winawer et al. 2013).

Critically, the association between these novel aperiodic metrics and cognitive states mirror the findings of studies using expensive and invasive techniques that derive E:I balance via gamma-aminobutyric acid (GABA) and glutamate expression (Gao et al. 2017). Due to the association of aperiodic metrics with multiple cognitive states, the potential for more accurate assessments of parameterized periodic activity, and cost-effective non-invasive measurements of E:I balance and activity akin to BOLD, novel parameterization techniques show promise and open new avenues of research in the fields of neuropathology and aging (Voytek and Knight 2015). However, their use as biomarkers assumes stability within individuals and minimal variance between individuals across time.

The test-retest reliability of total periodic activity is well established, with good to excellent intraclass correlation coefficients (ICC; Ding et al. 2022; Ip et al. 2018; McEvoy et al. 2000) over the scalp for eyes closed (EC) resting EEG (e.g., total theta, alpha, and beta power ICC = .84 - .97; Ip et al. 2018) and for eyes open (EO) resting EEG (e.g., total theta, alpha, and beta power ICC = .55 - .75; Ding et al. 2022). However, task-related EEG test-retest reliability is more varied and task-dependent. McEvoy et al. (2000) report good to excellent ICC during a working memory and psychomotor vigilance task (ICC >.7), while Ip et al. (2018) reported poor to excellent ICC for total theta (ICC = .37 - .83) and alpha power (ICC = .19 - .56) across four sessions of an auditory steady state response task.

Although few studies have examined the test-retest reliability of EEG metrics following parameterization of periodic and aperiodic activity (Levin et al. 2020; Pathania et al. 2021; Popov et al. 2023; Webb et al. 2023), they have also yielded inconsistent findings (Lopez et al. 2023). For example, while the short-term (∼ six days) reliability of resting aperiodic activity in children with Autism Spectrum Disorder (ASD) is moderate (aperiodic offset ICC = .53, aperiodic exponent ICC = .70), its reliability in typically developing (TD) children is poor to moderate (aperiodic offset ICC = .48, aperiodic exponent ICC = .70; Levin et al. 2020). This is also consistent across longer durations (i.e., 6 weeks; Webb et al. 2023). Furthermore, ICC for young adult resting-state recordings of the aperiodic exponent approximately 30 minutes apart are good to excellent (ICC’s .78 - .93) over frontal, central, parietal, and occipital regions of interest (ROI), and when using parameterization settings that limit non-neural artifact by excluding high beta and gamma activity (Pathania et al. 2021). This appears to be consistent after 1 week even in the presence of major drivers of between-subject variance, such as age (Popov et al. 2023). Nevertheless, other aspects of parameterized periodic activity (e.g., central frequencies of periodic activity) need to be considered as well as the reliability of the *SpecParam* model to fit across a nonrestricted frequency range.

### The current study

Identifying the conditions that yield optimal, or at least reliable, parameterized EEG activity recordings while also reducing the amount of data loss has received limited attention. Here, we conduct a secondary analysis of existing data previously reported in Wang et al. (2022a) and provide recommendation for the use and interpretation of parameterized data when implementing *SpecParam*. This is critical if *SpecParam* and similar techniques are to be used for large-scale evaluation of non-invasive measures relating to healthy aging and disease.

We use ICC to examine the test-retest reliability of parameterized activity across three sessions (90 minutes apart and 30 days later) in 49 healthy young adults at rest with EO and EC as well as during three EC cognitive tasks including subtraction (Math), music recall (Music), and episodic memory (Memory). In addition to ICC across each testing session, we also examine how consistent the ICC are across the scalp and identify the regions that provide the most, and the least, reliable measures of parameterized activity. Finally, we examine which conditions yield the most appropriate data for parameterization. Our results reveal that the test-retest reliability of parameterized periodic and aperiodic activity is generally good in EC and cognitive task-specific EEG. Conversely, the reliability of periodic and aperiodic activity in EO resting EEG performs poorly. In addition, the discrimination of theta peaks is poor following parameterization. Considering this, caution is needed when utilizing *SpecParam* to parameterize EO EEG, due to less reliable fits and the reduced presence of parameterized periodic peaks.

## METHODS

### Data availability

The processes and properties of the original data are described in full by Ding et al. (2022); Wang et al. (2022a). Processed data and scripts for the current study are available at https://github.com/MindSpaceLab/Aperiodic_Test_Retest.

### Participants

The original sample of 60 participants is publicly available at OpenNeuro (“Dataset ds004148”; https://openneuro.org/datasets/ds004148/versions/1.0.1; Wang et al. 2022b). Participants were aged between 18 and 28 (mean = 20.01 ± 1.88) and consisted of 32 females and 28 males. In accordance with the Review Board of the Institute of Southwest University, written and informed consent after a detailed explanation of the protocol was obtained from each participant. All experimental procedures were conducted in accordance with the Declaration of Helsinki. The Bond University Human Research Ethics Committee approved our secondary analyses of this data.

### Task description

Data collection occurred during three sessions across two lab visits. Following Session 1, participants were required to wait 90 minutes before completing Session 2 during their first lab visit. Participants returned to the lab approximately 30 days later (Session 3), with the visit occurring on the same day and time as their first visit. During each session, EEG data was recorded while participants completed two resting-state conditions (EO and EC) and three EC cognitive-state conditions (Math, Music, and Memory). For the EO resting-state recordings, participants were required to sit still, relax, and avoid blinking as much as reasonably possible. For the EC recordings, participants were required to stay awake. In each of the cognitive-state conditions, participants sat with their eyes closed and silently counted backwards from 5000 by 7s (Math), silently recalled and ‘sang’ their favorite song (Music) and recalled the events that had occurred that day from when they woke until their arrival at the laboratory (Memory). Participants performed the resting EO condition followed by the EC condition consecutively in each session before completing the cognitive-state conditions. Each cognitive-state condition was counterbalanced within each session but was consistent across the three visits. Each condition was five minutes long.

### Physiological recording and data reduction

EEG data was recorded from 63 Ag/AgCl active electrodes arranged according to the 10/20 system. Two electrode channels were used to record electrooculogram for eye movement detection while channel FCz was used as an online EEG reference. Signals were sampled at 500 Hz and resistance of all electrodes was kept below 5 kΩ. Preprocessing was conducted using EEGLAB (version 2019.1) functions and plugins implemented in MATLAB (version 2019b). Further detail on the preprocessing can be found in Ding et al. (2022). In our secondary analysis, the raw EEG data was re-referenced to a common average reference and was down sampled to 250 Hz and the PREP pipeline was implemented for further processing (Bigdely-Shamlo et al. 2015). Data was then filtered using an 8^th^ order Butterworth bandpass filter (1 – 45 Hz). Independent components analysis (ICA) in combination with Multiple Artifact Rejection Algorithm (MARA) was used to automatically identify and remove components associated with non-neural activity (e.g., eye movements, cardiac activity). Data from each condition was segmented into 2000ms epochs overlapping by 50%. Segments containing voltage changes of ±150 µV were rejected. We then used a Fast Fourier transformation (FFT; 2000ms Hamming window, 50% overlap, padded by a factor of 2) on the retained data from each participant to create a PSD for each channel within each condition and session. Eleven participants were excluded from further analyses as they had < 50% artifact-free data within any one condition or session (Table 1).

**Table 1.** Proportion of <50% artifact-free data in epochs for each participant across each Condition and Session.

| Session | ID | EC (%) | EO (%) | Math (%) | Memory (%) | Music (%) |
| --- | --- | --- | --- | --- | --- | --- |
| 1 | 11 | - | - | 65 | 62 | 61 |
|  | 13 | 79 | - | 94 | 97 | 87 |
|  | 23 | 71 | - | - | 73 | 67 |
|  | 47 | 61 |  | 65 | 61 | 62 |
|  | 58 | - | - | 68 | 77 | - |
| 2 | 05 | 73 | - | - | 61 | 63 |
|  | 13 | 90 | - | - | 72 | 89 |
|  | 23 | 86 | - | 87 | 67 | 70 |
|  | 58 | 78 | - | 81 | 86 | 71 |
| 3 | 13 | - | - | - | 67 | 55 |
|  | 17 | - | - | - | 55 | 80 |

*EC: Eyes closed; EO: Eyes open. Values are reported as percentage (%) of total epochs with artifact data.*

### Spectral parameterization

Prior to spectral parameterization of the PSD, total theta, alpha, and beta periodic activity was quantified using traditional methods of identifying the average power within canonical frequency bands (theta: 4 – 7 Hz; alpha: 8 – 13 Hz; beta: 14 – 30 Hz)^1^. PSD were imported into Python (version 3.9.16) for further spectral parameterization and statistical analysis.

Spectral parametrization was performed using the *SpecParam* package (https://github.com/fooof-tools/fooof; Donoghue et al. 2020b). For each PSD, we fit the *SpecParam* model in the semi-log space (logPower) between 2 and 40 Hz (peak width limits: 1-8; max number of peaks: 8; minimum peak height: 0.1; peak threshold: 2 SD; aperiodic mode: fixed) with a .25 Hz frequency resolution. For each channel, we extracted aperiodic parameters (exponent and offset), and parameterized periodic parameters for theta, alpha, and beta frequency bands. ANOVAs were performed with post hoc Tukey’s Honest Significant Difference (HSD) tests to determine if the number of unsuitable fits differed between each condition within each session. We then excluded participants with *SpecParam* model fits < 0.9 R^2^ on more than 50% of the channels across the three sessions within each condition. As such, the available sample size varied between conditions, but not between sessions. We opted for this approach as excluding participants across sessions and conditions resulted in a substantially reduced sample size due to poor fits over polar, temporal, and occipital electrodes in the EO condition. In analyses involving parameterized periodic activity, we excluded participants without identifiable peaks on > 50% of channels, applying this criterion across each session and within each condition. ICC were calculated for each channel using the Python port of the *irr* package (pyirr; de Klerk 2022; Gamer 2012). Similar to previous studies (Ding et al. 2022), we used the ICC calculation based on Shrout and Fleiss (1979) to determine the test-retest reliability of each periodic and aperiodic parameter. ICC analysis determines the ratio of intra-subject variability compared to all sources of variability for each individual repeated measure. ICC values of < .50 are considered poor, .50 to .75 are considered moderate, .75 to .90 are considered good, and > .90 are considered excellent^2^.

## RESULTS

### Data quality and *SpecParam* performance

Parameterization of EEG data from each participant using *SpecParam* generally performed well. Mean fit across the whole scalp (prior to exclusion of participant recordings <50% good fits) for each participant across each session and each condition was .95 ± .02 for EC, .92 ± .12 for EO, .95 ± .03 for Math, .95 ± .03 for Memory, and .95 ± .03 for Music. Results of the ANOVA assessing the mean model fit by condition indicated that there was a main effect of condition for Session 1 (*F_4, 275_* = 15.115, *p* < .001, η*²* = 0.18), Session 2 (*F_4, 275_* = 14.129, *p* < .001, η*²* = 0.17), and Session 3 (*F_4, 275_* = 7.885, *p* < 0.001, η*²* = 0.10). Tukey’s HSD post hoc analysis identified that model fit parameters were consistently poorer during EO recordings with significantly fewer good fits compared to all other conditions (Session 1: mean diff > 7.14, *p* < .001; Session 2: mean diff > 8.64, *p* < .001; Session 3: mean diff > 10.14, *p* < .0003). This appeared to primarily be over non-central sites (Figure 1A).

**Figure 1.**
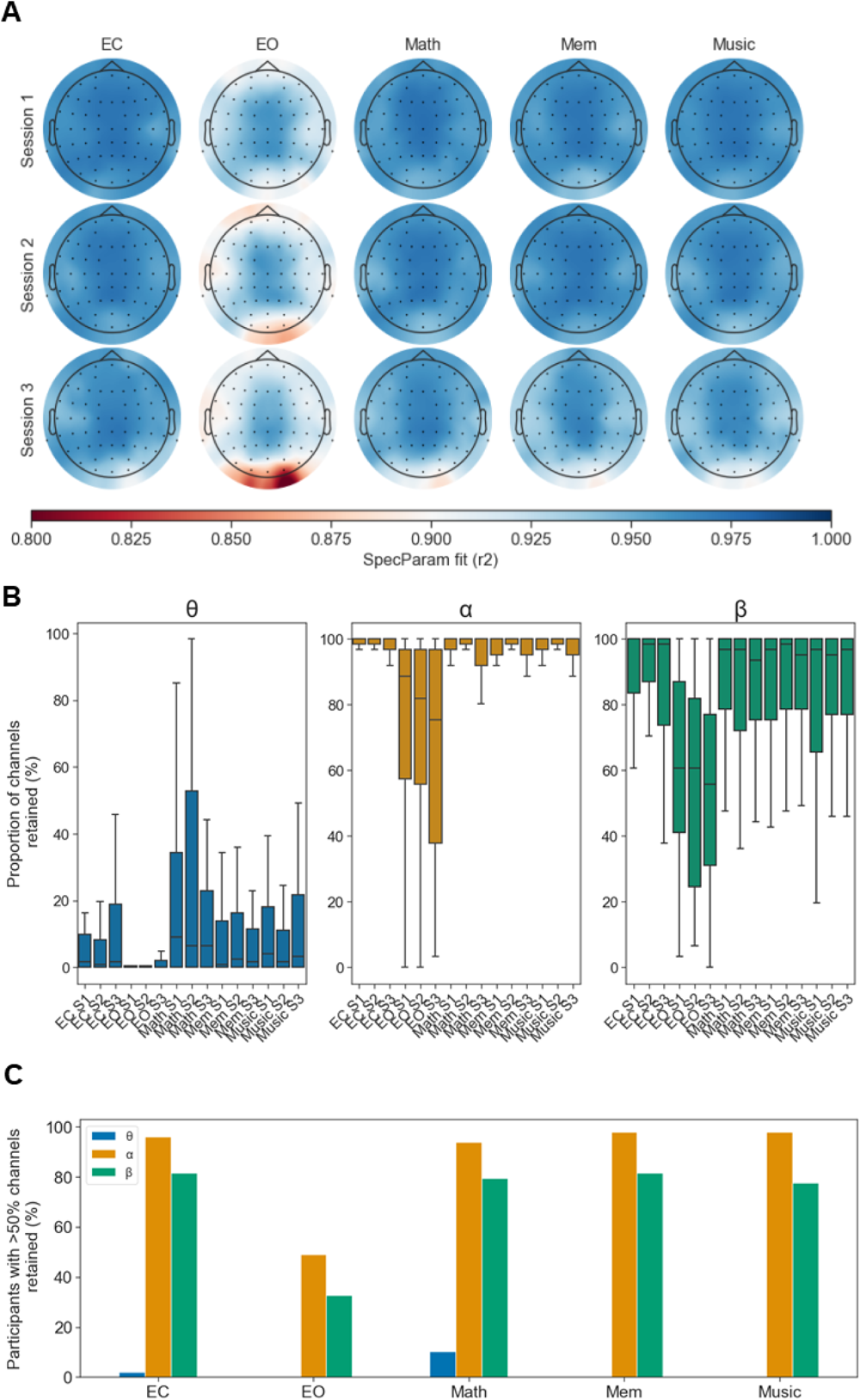
**Mean *SpecParam* model fit across the scalp (A), proportion of EEG channels retained for theta (θ), alpha (α) and beta (β) activity during each session of each condition (B), and proportion of participants who had > 50% of channels retained for each condition (C).** In general, parameterized θ activity was infrequently identified, leading to a lower number of retained channels in all conditions apart from the second session of Math. Similarly, EO had worse retainment of channels compared to EC, with most alpha and beta data loss coming from the EO sessions. Nonetheless, α and β generally had excellent retainment of channels across all conditions and sessions.

When assessing the performance of the *SpecParam* model fit across sessions for each condition, the ANOVA indicated that there were no significant differences in model fits between sessions for EC (*F_2, 165_* = 2.82, *p* = .062, η*²* = 0.03), EO (*F_2, 165_* = 2.87, *p* = .059, η*²* = 0.03), Math (*F_2, 165_* = 2.46, *p* = .088, η*²* = 0.03), or Music (*F_2, 165_* = 2.63, *p* = .074, η*²* = 0.03).

However, there was a significant difference in model fits across sessions for Memory (*F_2, 16_*

After the exclusion of participant recordings with < 50% good fits across all conditions and sessions, the sample size was decreased for Session 1 (*N =* 5 removed, 2.04%), Session 2 (*N* = 10 removed, 4.08%) and Session 3 (*N* = 27 removed, 11.02%). Although there were some instances of poor fits in other conditions (Table 2), these were not as profound as resting EO data (14.96% of total recordings across the three sessions removed). We also detected variance in the retainment of parameterized oscillatory peaks between theta, alpha, and beta bands across conditions and sessions (Figure 1B). In each condition (Figure 1C), theta band activity was poorly retained, with only one (1.8%) and five (9.25%) participants having detectable theta peaks in > 50% of channels, and only in the EC and Math conditions, respectively. Indeed, theta was only detected in .0001% of all fits (e.g., participant × session × condition × channel). Alpha band activity was excellently retained for EC (*N* = 47, 95.91%), Math (*N* = 46, 93.87%), Memory and Music (*N* = 48, 97.95%), but poorly for EO (*N* = 24, 48.97%). The retainment of beta band activity was moderate to good for EC (*N* = 40, 81.63%), Math (*N* = 39, 79.59%), Memory (*N* = 40, 81.63%), Music (*N* = 38, 77.55%), but poor for EO (*N* = 16, 32.65%).

**Table 2.** Participant recordings with poor SpecParam model fits for within conditions and within sessions.

| Condition | Session |  |  | Total |
| --- | --- | --- | --- | --- |
|  | 1 | 2 | 3 |  |
| EC | - | 2 (4.08) | 3 (6.12) | 5 (3.40) |
| EO | 4 (8.16) | 6 (12.24) | 12 (24.48) | 22 (14.96) |
| Math | 1 (2.04) | 1 (2.04) | 4 (8.16) | 6 (4.08) |
| Memory | - | - | 4 (8.16) | 4 (2.22) |
| Music | - | 1 (2.04) | 4 (8.16) | 5 (3.40) |
| Total | 5 (2.04) | 10 (4.08) | 27 (11.02) |  |

*EC: Eyes closed condition; EO: Eyes open condition. Values are reported as N (%) of participant recordings out of a total of 147 recordings for within conditions and 245 recordings for within sessions.*

In summary, the general performance of alpha and beta activity was moderate to good, with minimal data loss in EC, Math, Memory, and Music. In contrast, most participants did not have clear parameterized alpha or beta peaks in > 50% of channels in the EO condition. Owing to the substantially reduced sample size due to bad fits in non-central sites in EO recordings, subsequent analyses adopted a pair-wise approach to exclusions.

### Test-retest reliability

Test-retest reliability was assessed for total theta, alpha, and beta power prior to parameterization (Figure 2A). Consistent with previous studies (Table 3; Ding et al. 2022), total theta had good reliability for EC (mean ICC = .85), EO (mean ICC = .83), Math (mean ICC = .89), Memory (mean ICC = .82), and Music (mean ICC = .86). Similarly, total alpha had good reliability for EC (mean ICC = 0.83), EO (mean ICC = .89), Math (mean ICC = .88), Memory (mean ICC = .89), and Music (mean ICC = .87). Total beta activity generally had the lowest reliability, but this was good for EC (mean ICC = .76), EO (mean ICC = .82), Math (mean ICC = .82), Memory (mean ICC = .84), and Music (mean ICC = .83; Figure 2B).

**Figure 2.**
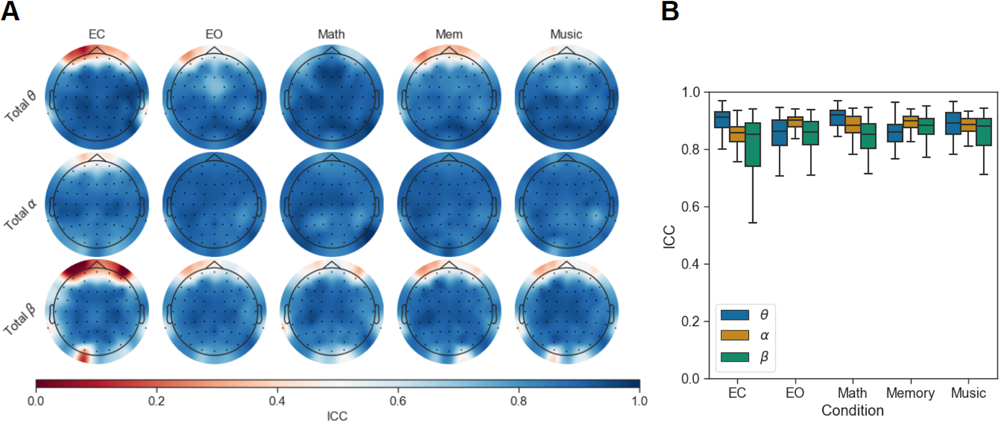
**Intraclass correlation coefficients for total periodic band power before *SpecParam* parameterization of EEG.** In general, total periodic theta (θ), alpha (α), and beta (β) activity had high test-retest reliability. However, total beta band activity had the poorest reliability compared to the other periodic bands. Boxplots indicate the mean, IQR, and range of ICC across electrodes.

**Table 3.** Average ICC for each EEG measure across the eye and cognitive task conditions.

|  | EC | EO | Math | Memory | Music |
| --- | --- | --- | --- | --- | --- |
| Total theta power | .85 | .83 | .89 | .82 | .86 |
| Total alpha power | .83 | .89 | .88 | .89 | .87 |
| Total beta power | .76 | .82 | .82 | .84 | .83 |
| Parameterized alpha power | .83 | .79 | .81 | .82 | .83 |
| Parameterized alpha CF | .91 | .85 | .91 | .92 | .93 |
| Parameterized alpha BW | .63 | .58 | .63 | .68 | .77 |
| Parameterized beta power | .79 | .62 | .77 | .81 | .78 |
| Parameterized beta CF | .81 | .79 | .80 | .85 | .82 |
| Parameterized beta BW | .65 | .55 | .72 | .67 | .70 |
| Exponent | .73 | .70 | .71 | .64 | .70 |
| Offset | .85 | .81 | .85 | .81 | .84 |

*ICC: Intraclass correlation coefficient; EEG: electroencephalography; EC: Eyes closed condition; EO: Eyes closed condition; CF: Central frequency; BW: Bandwidth*

Test-retest reliability was then assessed for parameterized alpha and beta activity, once isolating periodic activity from aperiodic activity (Figure 3A). Theta activity was excluded as there was only distinguishable theta activity in five participants (Figure S2 PSYCHOMETRIC ANALYSIS OF SPECTRAL PARAMETERIZATION of supplementary material). Parameterized alpha power and central frequency were generally good to excellent during EC (mean ICC = .83 and .91), EO (mean ICC = .79 and .85), Math (mean ICC = .81 and .91), Memory (mean ICC = .82 and .92) and Music (mean ICC = .83 and .93). Parameterized alpha bandwidth performed the poorest, with moderate to good ICC for EC (mean ICC = .63), EO (mean ICC = .58), Math (mean ICC = .63), Memory (mean ICC = .68), and Music (mean ICC = .77; Figure 3B). Parameterized beta power and central frequency was good but did not perform as well as parameterized alpha for EC (mean ICC = .79 and .81), EO (mean ICC = .62 and .79), Math (mean ICC = .77 and .80), Memory (mean ICC = .81 and .85), and Music (mean ICC = .78 and .82). Parameterized beta bandwidth performed the poorest, with moderate to good ICCs for EC (mean ICC = .65), EO (mean ICC = .55), Math (mean ICC = .72), Memory (mean ICC = .67), and Music (mean ICC = .70; Figure 3C). Considering this, test-retest reliability of parameterized periodic activity performs best over frontocentral and parietal sites and is worst over non-central sites. This was similar across conditions, data loss notwithstanding.

**Figure 3.**
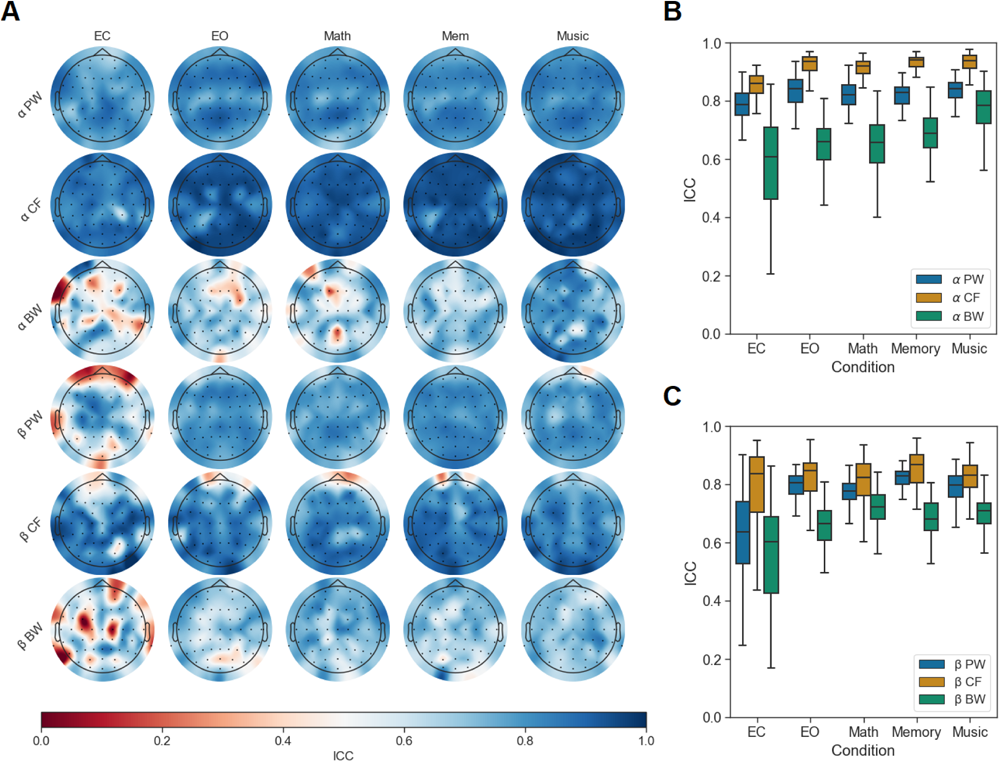
Intraclass correlation coefficients for parameterized periodic band activity following *SpecParam* parameterization of EEG. In general, parameterized alpha (α) had excellent while parameterized beta (β) had good test-retest reliability. In both the parameterized alpha and beta bands, bandwidth performed the poorest. Boxplots indicate the mean, IQR, and range of ICC across electrodes.

Test-retest reliability was also assessed for aperiodic measures (Figure 4A). The aperiodic exponent generally had good test-retest reliability for EC (mean ICC = .73), EO (mean ICC = .70), Math (mean ICC = .71), and Music (mean ICC = .70), but moderate for Memory (mean ICC = .64). The aperiodic offset performed substantially better with good reliability for EC (mean ICC = .85), EO (mean ICC = .81), Math (mean ICC = .85), Memory (mean ICC = .81) and Music (mean ICC = .84; Figure 4B).

**Figure 4.**
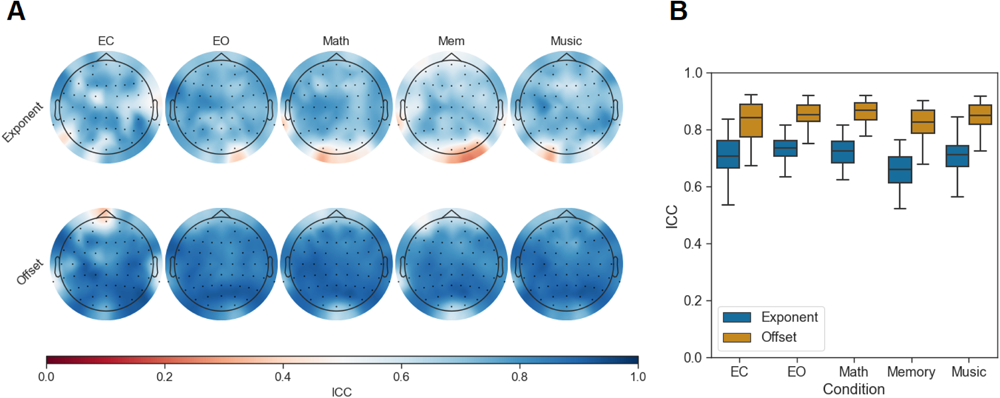
Intraclass correlation coefficients for aperiodic exponent and offset following *SpecParam* parameterization of EEG. Although both the aperiodic exponent and offset demonstrated good test-retest reliability in both eye and task conditions, the aperiodic offset outperformed the aperiodic exponent in all eye and task conditions. Boxplots indicate the mean, IQR, and range of ICC across electrodes.

## DISCUSSION

In general, the *SpecParam* technique for assessing aperiodic activity of EEG in healthy young adults is consistent across conditions and stable across time in most participants and states. However, EO resting data is problematic due to the *SpecParam* technique producing poor fits. The current study aimed to assess the test-retest reliability of total periodic activity, parameterized periodic activity, and aperiodic activity in a population of healthy young adults across 3 repeated sessions where EEG was collected during resting states and cognitive tasks. When following a *SpecParam* model fit inclusion threshold of > .90 R^2^ in at least 50% of EEG channels, nearly a third of participants were lost due to inadequate fits, and this was primarily seen in the EO condition. However, for retained participants *SpecParam* model fitting performed generally well for alpha and beta activity in all states, excluding EO, but was only able to distinguish theta band activity in < 5 people. The test-retest reliability for total and parameterized alpha activity was good to excellent, while total and parameterized beta activity was good. Finally, the aperiodic exponent, and to a greater extent the aperiodic offset, demonstrated good test-retest reliability.

### Performance of *SpecParam* parameterization

The recent development of algorithms to parameterize EEG has enabled the refined assessment of periodic and aperiodic activity during resting and task related states (Donoghue et al. 2020b; Kosciessa et al. 2020; Wen and Liu 2016). Although this has been widely beneficial for understanding underlying mechanisms of E:I balance in neural networks, the sensitivity of *SpecParam* to identify and accurately fit a wide range of periodic activity has yet to be assessed. In line with Donoghue et al. (2020b), we have demonstrated that *SpecParam* performs excellently for fitting alpha and beta peaks. However, we found that its ability to identify slower theta activity and during EO states, is questionable.

Once *SpecParam* model fitting was undertaken, only 5 participants had identifiable theta activity in only the EC and Math conditions. Although theta activity did increase during the Math task, this was only marginally different to the resting EC state. Previous evidence suggests that theta at rest is minimal and may be dependent on cognitive task performance (Cavanagh and Frank 2014). Indeed, the presence of theta and its power once parameterized is more variable compared to other oscillatory bands (Donoghue et al. 2020b). In a recent assessment of theta-beta ratios in younger and older adults (Finley et al. 2022), only 15% of participants had definable parameterized fronto-central theta peak following *SpecParam* model fitting. Similar findings have also been seen when looking at a smaller age range (50 – 80 years) where theta power is unidentifiable (Smith et al. 2023). In a similar assessment of theta reliability to the current study, theta activity was indistinguishable above the aperiodic component after utilizing parameterization of the PSD, particularly during resting EC states (Popov et al. 2023). Considering this, the influence of theta activity once *SpecParam* model fitting has occurred should be interpreted with caution, as it may be an unknown combination of aperiodic and periodic activity.

Surprisingly, we observed that the retainment of periodic activity in the PSD of each channel was poorest during the EO condition, particularly over non-central electrode sites. On the contrary to our study of young adults, Hill et al. (2022) observed that the *SpecParam* model fit is best during EO (R^2^ = .99; Error = .07) compared to EC (R^2^ = .98; Error = .05) in TD children (age range: 4-12 years). However, beyond this, it is unclear how eye state influences *SpecParam* model fitting. As alpha power and alpha dynamics are most prominent when eyes are closed (Donoghue et al. 2020a), periodic peaks may have been more easily distinguishable during the EC compared to the EO state in the current study. Nonetheless, as the model fit represents the average across the whole scalp, special consideration needs to be given regarding ROI electrode selection.

### Test-retest reliability of aperiodic and periodic activity

The test-retest reliability of aperiodic activity in the current study was moderate to good for the exponent, and good for the offset. Furthermore, the offset was substantially more stable and consistent than the exponent. Previous investigations of the test-retest reliability of aperiodic activity have been inconclusive. In cohorts of TD children (Levin et al. 2020; Webb et al. 2023), aperiodic activity has poor to moderate reliability. However, this may be due to neurodevelopment occurring more dynamically in TD children compared to young adults. Indeed, when assessed in a young adult cohort comparable to that of the current study (Pathania et al. 2021), the test-retest reliability of the exponent and offset are considered good to excellent over frontal, central, parietal, and occipital ROI.

Previous assessments of test-retest reliability of total periodic activity have demonstrated that the stability and consistency of theta, alpha, and beta activity is excellent during resting and task-related tasks. Furthermore, the test-retest reliability of traditional EEG band power metrics have not distinguished differences between EC and EO data (Pollock et al. 1991). In the current study, we build upon this evidence by elucidating the test-retest reliability of parameterized periodic activity once it has been isolated from underlying aperiodic activity. Consistent with previous investigations of total periodic activity using EEG (Ding et al. 2022; Ip et al. 2018; Levin et al. 2020; McEvoy et al. 2000; Pathania et al. 2021; Popov et al. 2023; Webb et al. 2023), we found the test-retest reliability of total theta, alpha, and beta activity to be good during rest and cognitive-states. Total beta activity had the poorest reliability and had the greatest amount of variability in ICC, especially around non-central sites during EC and the Math conditions. As theta and alpha band activity is related to global neural processing across the whole brain whereas beta activity is related to local neural processing at specific brain regions (Ding et al. 2022; Martin-Buro et al. 2016), the confounding influence of aperiodic activity may be greatest for beta test-retest reliability compared to other periodic bands when assessed scalp-wide. This highlights the importance of accounting for aperiodic activity in the EEG.

Following parameterization, reliability of parameterized alpha power and central frequency was generally good to excellent, while parameterized beta power and central frequency was generally good. Due to the lack of detectable parameterized theta peaks, we are unable to comment on the test-retest reliability of parameterized theta activity. Nonetheless, our findings signify that parameterized power and frequency can be a reliable way of identifying individual differences at rest and during cognitive states. Although our findings relating to parameterized alpha and beta activity are to be expected (Levin et al. 2020; Pathania et al. 2021; Webb et al. 2023), the significantly poorer performance of parameterized alpha and parameterized beta bandwidth scalp-wide was not anticipated. Of the previous investigations of test-retest reliability of parameterized periodic activity, Levin et al. (2020) found a decrease in the parameterized alpha bandwidth ICC when adjusting for age in TD children, which was suggested to be due to increasing alpha peak frequencies across the six days of investigation. Although this coincides with the findings of the current study, why bandwidth ICC were poorer than the other parameters of periodic activity, and what this signifies, is unclear.

The stability and consistency of the aperiodic exponent and offset supports the notion of using aperiodic activity as a neural biomarker, even in samples that are homogenous with respect to a major driver of between-subject variance (i.e., the confounding effect of age; Finley et al. 2022; Voytek et al. 2015). Furthermore, areas that generally have weaker oscillations will perform worse from a data quality standpoint, and these tend to be non-central areas. However, with retained participants, test-retest reliability of periodic and aperiodic activity is generally good.

### Considerations

We have intentionally used a stringent inclusion criterion preceding the test-retest reliability analysis. Unlike previous studies that have only focused on a small cluster of electrodes (i.e., electrodes centralised around Pz or Fz; Levin et al. 2020; Pathania et al. 2021) that typically yield strong oscillatory activity, we excluded participants based on poor fits/missing peaks on areas and in conditions that are systematically worse. Regarding the absence of theta following parameterization, it is possible that either *SpecParam* was unable to detect theta peaks in the PSD, the tasks were not overly cognitively demanding, or distinguishable theta peaks were not present initially. Considering theta activity is minimal at rest and increases with cognitive demand (Cavanagh and Frank 2014), it is likely that a combination of suboptimal performance of *SpecParam* in fitting theta oscillations, as well as indistinguishable peaks in the PSD, contributed to the exclusion of parameterized theta activity. Because of this, the inability of *SpecParam* to resolve this theta issue needs further consideration.

Recently, the role of aperiodic constituents of event-related potentials in the EEG during sensory, motor, and cognitive events has gained interest (Arnett et al. 2022; Cadwallader et al. 2023; Virtue-Griffiths et al. 2022). Although the findings of the current study provide insight into the test-retest reliability of aperiodic measures during relatively simple cognitive tasks, these findings are distinct to cognitive tasks that are more complex (i.e., working memory task). Considering this, future work needs to identify the test-retest reliability of aperiodic activity during more complex cognitive tasks that consider time-locked events.

Another consideration is the duration of EEG data implemented in *SpecParam* and if parameterized theta activity would have been present if a longer duration of data was used. Previous research has investigated the performance of *SpecParam* using split-half time to examine within-session consistency of EEG activity in TD children and ADHD adolescents across the whole scalp (Karalunas et al. 2022). *SpecParam* performed well even with < 1 minute of data included. Considering this, future research may benefit from including longer durations of EEG data, larger datasets, and by also comparing the performance of *SpecParam* to other models used to distinguish between periodic and aperiodic activity.

## Conclusion

Although tools such as *SpecParam* allow for the development of new biomarkers based on parameterized neural activity, few studies have assessed the test-retest reliability of parameterized measures. This study indicates that *SpecParam* is a promising tool for parameterizing periodic activity from aperiodic activity and is appropriate for identifying alpha and beta peaks in some states and scalp locations. Although this study demonstrates that the stability and consistency of parameterized periodic and aperiodic activity is good, careful consideration needs to be taken when utilizing *SpecParam* in resting EO states. That is, EO states may result in poor *SpecParam* model fitting. Furthermore, data quality with regards to fit, detection/presence, and research goal, needs to be considered as *SpecParam* may not be appropriate for identifying slow oscillatory activity, such as theta. Overall, it appears that the aperiodic exponent and offset have the potential for being a reliable and consistent biomarker.

## ACKNOWLEDGEMENTS

We would like to express our gratitude to all the participants involved in our study, for their support and patience, and for contributing their time.

## DATA AND CODE AVAILABILITY STATEMENT

The raw data used in this study is available at OpenNeuro (“Dataset ds004148”; https://openneuro.org/datasets/ds004148/versions/1.0.1). Processed data and for all analyses and plots are publicly available at https://github.com/MindSpaceLab/Aperiodic_Test_Retest.

## AUTHOR CONTRIBUTIONS

Conceptualization: A.J.F., N.K., J.F.C., H.A.D.K, and D.J.A. Data curation: D.J.M., J.F.C., and D.J.A. Formal analysis: D.J.M. and D.J.A. Project administration: J.F.C. and D.J.A. Visualization: D.J.A. Writing - original draft: D.J.M. and D.J.A. Writing - review & editing: D.J.M., A.J.F., N.K., O.B., V.S., A.A.M., J.F.C., H.A.D.K, and D.J.A.

## Footnotes

1 For quantification of total delta and gamma periodic activity of the current dataset see Ding et al. (2022).

2 ICC values for aperiodic activity were also examined with the *IRASA* tool which found similar but smaller results (see Figure S1 and Table S1 of supplementary material). = 3.31, *p* = .038, η*²* = 0.03). Tukey’s HSD post hoc analysis identified that the model fit was significantly worse at Session 3 compared to Session 2 (mean diff = 3.78, *p* = .039) but not Session 1 (mean diff = 2.91, *p* = .14). Furthermore, there was no difference in model fit between Session 1 and Session 2 (mean diff = -.87, *p* = .83).

