## Supplementary files for "Test-retest reliability of spectral parameterization by 1/*f* characterization using *SpecParam*"

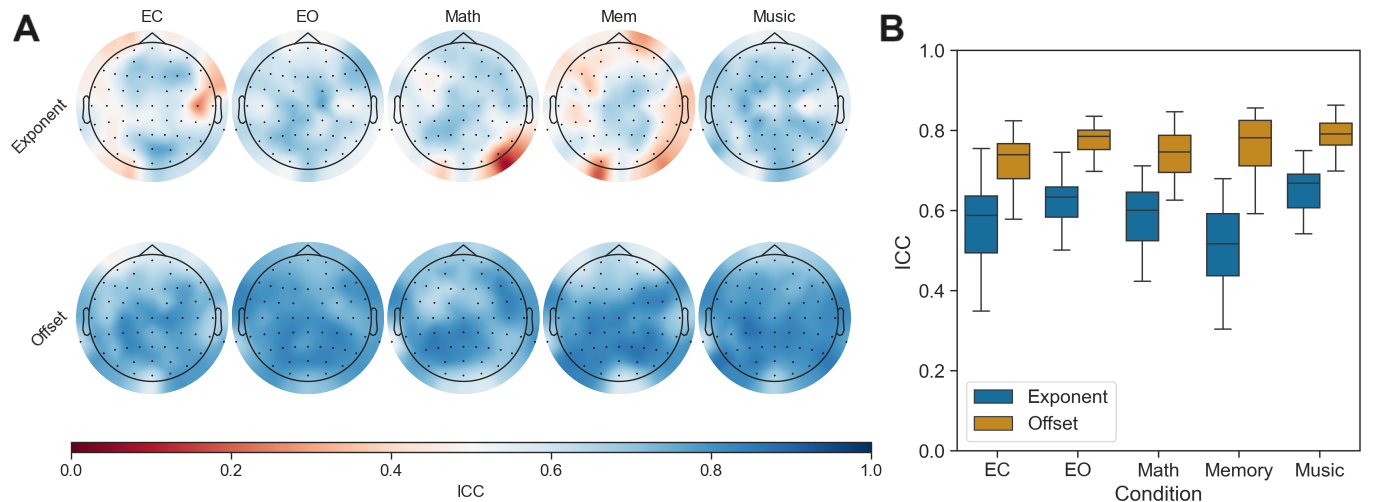

**Figure S1. Intraclass correlation coefficients for aperiodic exponent and offset following *IRASA* parameterization of EEG.** Similar to *SpecParam,* both the aperiodic exponent and offset demonstrated good test-retest reliability in both eye and task conditions, the aperiodic offset outperformed the aperiodic exponent in all eye and task conditions. Boxplots indicate the mean, IQR, and range of ICC across electrodes.

*Table S1.* ICCs comparison between *IRASA* and *SpecParam* tools for parameterising aperiodic activity.

|  | *IRASA* | | | | | *SpecParam* | | | | |
| --- | --- | --- | --- | --- | --- | --- | --- | --- | --- | --- |
|  | EC | EO | Math | Mem | Music | EC | EO | Math | Mem | Music |
| Exponent | .63 | .57 | .58 | .51 | .65 | .73 | .70 | .71 | .64 | .70 |
| Offset | .78 | .72 | .74 | .76 | .78 | .85 | .81 | .85 | .81 | .84 |

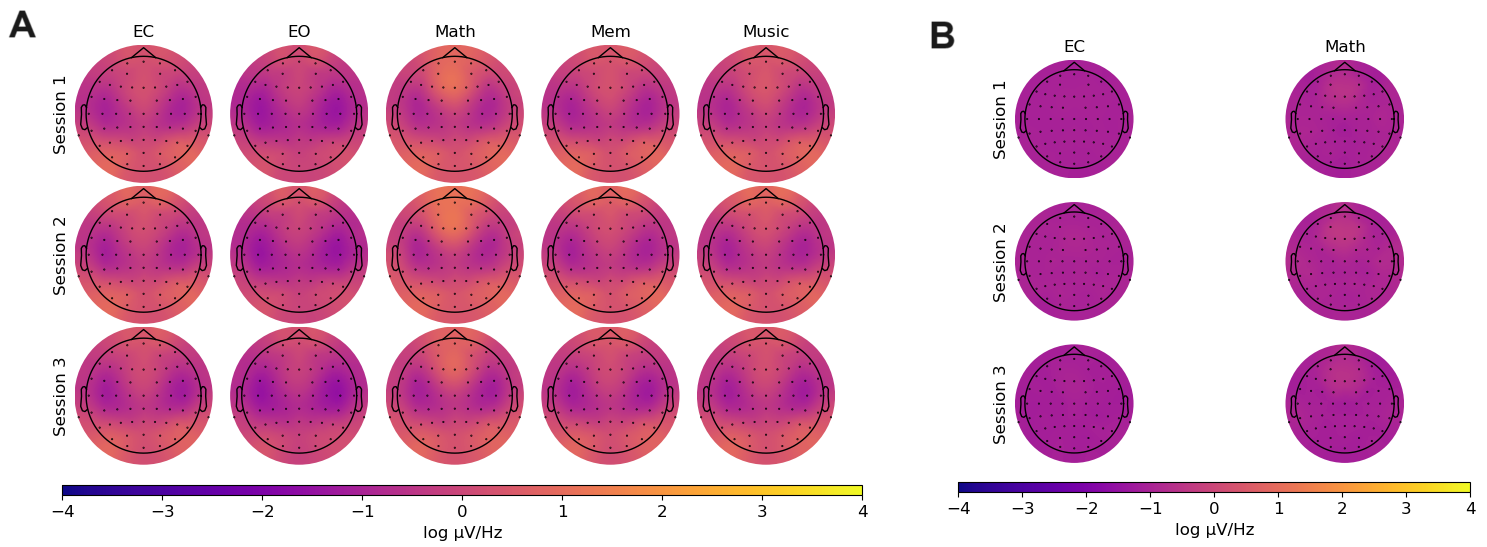

**Figure S2. Theta power prior (A) and following (B) spectral parameterization using *SpecParam*.** Theta activity was primarily elicited during the Math condition. However, following parameterization, only 1 participant and 5 participants for the EC and Math conditions respectively had detectable theta peaks >50% of channels.
